# Characterization of a glycolipid glycosyltransferase with broad substrate specificity from the marine bacterium *Candidatus* Pelagibacter sp. HTCC7211

**DOI:** 10.1101/2021.03.25.437072

**Authors:** Tao Wei, Caimeng Zhao, Mussa Quareshy, Nan Wu, Shen Huang, Yuezhe Zhao, Pengfei Yang, Duobin Mao, Yin Chen

## Abstract

In the marine environment, phosphorus availability significantly affects the lipid composition in many cosmopolitan marine heterotrophic bacteria, including members of the SAR11 clade and the Roseobacter clade. Under phosphorus stress conditions, non-phosphorus sugar-containing glycoglycerolipids are substitutes for phospholipids in these bacteria. Although these glycoglycerolipids play an important role as surrogates for phospholipids under phosphate deprivation, glycoglycerolipid synthases in marine microbes are poorly studied. In the present study, we biochemically characterized a glycolipid glycosyltransferase (GT_cp_) from the marine bacterium *Candidatus* Pelagibacter sp. HTCC7211, a member of the SAR11 clade. Our results showed that GT_cp_ is able to act as a multifunctional enzyme by synthesizing different glycoglycerolipids with UDP-glucose, UDP-galactose, or UDP-glucuronic acid as sugar donors and diacylglycerol as the acceptor. Analyses of enzyme kinetic parameters demonstrated that Mg^2+^ notably changes the enzyme’s affinity for UDP-glucose, which improves its catalytic efficiency. Homology modelling and mutational analyses revealed binding sites for the sugar donor and the diacylglycerol lipid acceptor, which provided insights into the retaining mechanism of GT_cp_ with its GT-B fold. A phylogenetic analysis showed that GT_cp_ and its homologs form a group in the GT4 glycosyltransferase family. These results not only provide new insights into the glycoglycerolipid synthesis mechanism in lipid remodelling, but also describe an efficient enzymatic tool for future synthesis of bioactive molecules.

**Importance:** The bilayer formed by membrane lipids serves as the containment unit for living microbial cells. In the marine environment, it has been firmly established that phytoplankton and heterotrophic bacteria can substitute phospholipids with non-phosphorus sugar-containing glycoglycerolipids in response to phosphorus limitation. However, little is known about how these glycoglycerolipids are synthesized. Here, we determined the biochemical characteristics of a glycolipid glycosyltransferase (GT_cp_) from the marine bacterium *Candidatus* Pelagibacter sp. HTCC7211. GT_cp_ and its homologs form a group in the GT4 glycosyltransferase family, and can synthesize neutral glycolipids (MGlc-DAG and MGal-DAG) and an acidic glycolipid (MGlcA-DAG). We also uncover the key residues for DAG-binding through molecular docking, site-direct mutagenesis and subsequent enzyme activity assays. Our data provide new insights into the glycoglycerolipid synthesis mechanism in lipid remodelling.

## Introduction

Phospholipids form the structural basis of all cells, but sugar-containing glycoglycerolipids are mainly restricted to marine microbes, cyanobacteria, and higher plants (1, 2). Glycoglycerolipids are found on the lipid bilayer of cell membranes and play critical roles in cell growth, cellular recognition, adhesion, neuronal repair, and signal transduction. These natural glycoglycerolipids often have unusual and sometimes unexpected biological activities, such as antitumor, antiviral, anti-inflammatory, antimalarial, immunostimulatory, and neuritogenic activities, which make them valuable molecular targets for research (3–5). The basic structure of glycoglycerolipids is characterized by a 1, 2-diacyl-*sn*-glycerol (DAG) moiety with different numbers and types of sugars (glucose, galactose, mannose, rhamnose, or charged sugars like glucuronic acid or sulfoquinovose) attached at the *sn*-3 position of the glycerol backbone in DAG. These sugar attachments have an α- or β-anomeric configuration, and are bound via (1→2), (1→3), (1→4), or (1→6) linkages (6, 7). The common glycoglycerolipid structures in marine heterotrophic microbes and cyanobacteria are 1,2-diacyl-3-O-(β-D-galactopyranosyl)-*sn*-glycerol (monogalactosyl diacylglycerol, MGal-DAG), 1,2-diacyl-3-O-(α-D-glucopyranosyl)-*sn*-glycerol (monoglucosyl DAG, MGlc-DAG),1,2-diacyl-3-O-(α-D-galactopyranosyl)-(1→6)-O-β-D-galactopyranosyl)-*sn*-glycerol (digalactosyl DAG, DGal-DAG), and 1,2-diacyl-3-O-(6-deoxy-6-sulfo-α-D-galactopyranosyl)-*sn*-glycerol (sulfoquinovosyl DAG, SQDG) (2,8).

Glycoglycerolipids are usually synthesized by glycosyltransferases (GTs), which are highly divergent and polyphyletic. The GTs can be categorized into 110 numbered families according to their sequence similarity and signature motifs, and the stereochemistry of the glycoside linkage formed (9). Of the 110 families, the families GT4, GT21, and GT28, which are known as glycoglycerolipid synthases, utilize sugar nucleotides as donors and contain a consensus sugar donor binding domain near the C-terminus (10, 11). Despite the wide variety of bacterial glycoglycerolipids, only a few bacterial lipid GTs have been identified and characterized so far. The GTs synthesizing MGlc-DAG and DGal-DAG have been isolated from the cell wall-less bacterium *Acholeplasma laidlawii*, and were found to belong to the GT4 family in the carbohydrate-active enzymes (CAZy) database (12). Other known members of bacterial GT4 include the MGlc-DAG synthases from *Deinococcus radiodurans* and *Thermotoga maritima* and the MGal-DAG synthase from *Borrelia burgdorferi*, which was the first cloned galactosyltransferase forming MGal-DAG with the α-anomeric configuration of the sugar (13, 14). A bifunctional GT (designated as Agt) from *Agrobacterium tumefaciens* was found to synthesize MGlc-DAG or MGlcA-DAG with UDP-glucose (UDP-Glc) or UDP-glucuronic acid (UDP-GlcA) as the sugar donor, respectively (7). This enzyme also belongs to the GT4 family and was the first glucuronosyl DAG synthase to be isolated. The processive GTs (Pgts), however, are members of the GT21 family, and show high sequence similarity to GT4 family GTs. The Pgts include enzymes from *Mesorhizobium loti* and *A. tumefaciens* that synthesize DGal-DAG, glucosylgalactosyl-DAG (GlcGal-DAG), and triglycosyl DAGs (15, 16). To the best of our knowledge, the structure function relationship of DAG-dependent GT4 glycosyltransferases has not been studied previously due to the lack of crystal structure of these glycoglycerolipids-producing enzymes and, as such, the binding pockets for UDP-sugars and DAG remain elusive.

Glycoglycerolipids play important roles in marine phytoplankton and heterotrophic bacteria under phosphate deprivation. Lipid remodelling reduces the cellular requirement for phosphorus, and the glycoglycerolipids MGlc-DAG/MGlcA-DAG and SQDG replace phospholipids in marine heterotrophic bacteria and marine phytoplankton and cyanobacteria, respectively (17–20). However, little is known about how these glycoglycerolipids are synthesized. Previous studies have shown that a manganese-dependent metallophosphoesterase, PlcP, is essential for lipid remodelling in marine heterotrophs, and that the *plcP* gene is organized in an operon-like structure and a putative glycosyltransferase was found down stream of *plcP* in numerous marine heterotrophic bacteria, such as members of the SAR11 clade (18, 21). Our previous work has shown that the GT from the marine bacterium SAR11 (*Candidatus* Pelagibacter sp. HTCC7211 and HTCC1062) is homologous to the Agt GT in *A. tumefaciens* (18). However, the activity of a SAR11 GT in the synthesis of glycoglycerolipids has not been characterized so far.

In this study, we report the detailed biochemical characterization of a glycoglycerolipid GT (GT_cp_) from the marine bacterium *Candidatus* Pelagibacter sp. HTCC7211. Our results showed that GT_cp_ has a broad substrate specificity and can synthesize neutral glycolipids (MGlc-DAG and MGal-DAG) and an acidic glycolipid (MGlcA-DAG). GT_cp_ represents the first member of the GT4 family of lipid GTs from marine bacteria. In addition, homology modelling and site-directed mutagenesis analyses revealed details of its substrate recognition mechanism and identified key residues involved in the co-ordination of DAG in a GT4 family glycosyltransferase for lipid biosynthesis.

## Results and discussion

### GT_cp_ and its homologs form a new group in the GT4 family with a GT-B fold structure

The gene encoding a putative GT_cp_ (WP_008545403.1) consists of 1005 bp encoding a peptide of 334 amino acids, containing a GT4-like domain (Fig. 1A). This gene was previously hypothesized to be involved in lipid remodelling in *Candidatus* Pelagibacter sp. HTCC7211 for the synthesis of MGlc-DAG and MGlcA-DAG (17, 18). Sequence alignment (Fig. 1B) analyses showed that the GT_cp_ amino acid sequence has between 46-58%sequence identity with the putative GT from *Labrenzia aggregata(GT_la_*, WP_040439323.1), *Thalassospira lucentensis* (GT_tl_, WP_062950653.1), *Methylophaga nitratireducenticrescens* (GT_mn_, WP_014706011.1), *Desulfobulbus mediterraneus* (GT_dm_, WP_028584068.1), *Citromicrobium bathyomarinum* JL354 (GT_cb_, WP_010239457.1), *Kordiimonas gwangyangensis* (GT_kg_, WP_051078133.1), and the characterized Agt (locus tag, atu2297) from *A. tumefaciens* (7). In the phylogenetic analysis, the GT_cp_, together with its close homologs (GT_la_, GT_tl_, GT_mn_, GT_dm_, GT_cb_, GT_kg_ and Agt), formed a clade in the GT4 family (Fig. 1A). Sequences from this clade showed low sequence identity (< 25%) to other members of GT4 family, which includes more than 150,000 proteins with at least 22 different enzymatic activities at the time of writing. Purified Agt from *A. tumefaciens* has been found to synthesize MGlc-DAG or MGlcA-DAG with UDP-glucose or UDP-glucuronic acid as the sugar donor, respectively, and the expression of *agt* is known to be induced under phosphate deficiency (18). Neither GT_cp_ nor any of its homologs from marine bacteria have been purified nor characterized to date.

**Figure 1.**
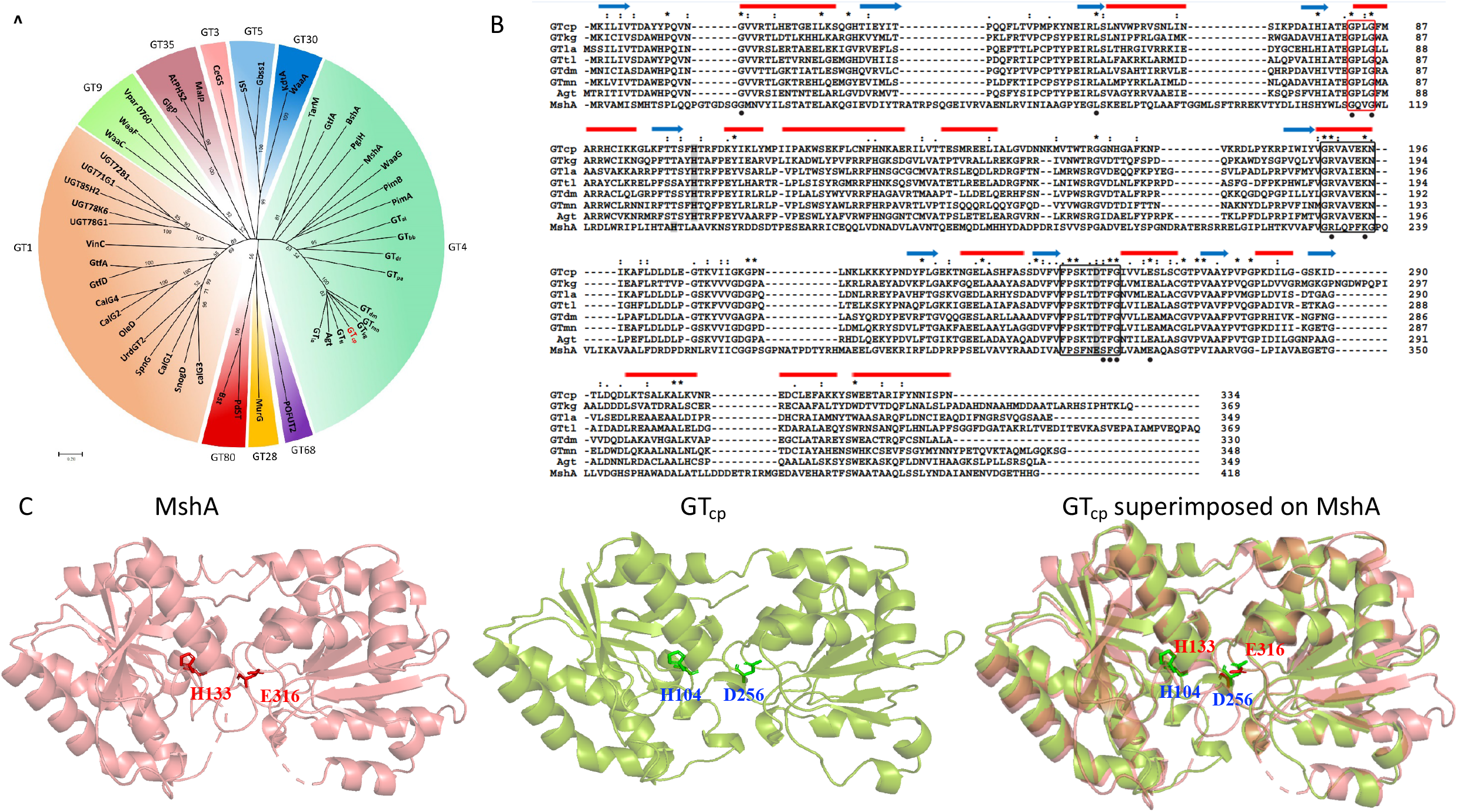
Multiple sequence alignment and functional domain analyses of GT_cp_ protein. (A) A phylogenetic tree of GT_cp_ and its homologs with known 3-dimentionalX-ray structure of GTs. Sequences and structures of GTs are obtained from the NCBI database and the PDB database, including GT1 (calG3, SnogD, CalG1, SpnG, UrdGT2, OleD, CalG2, CalG4, GtfD, GtfA, Vinc, UGT78G1, UGT78K6, UGT85H2, UGT71G1 and UGT72B1), GT3 (CeGs), GT5(Gbss1 and SSI), GT9 (WaaC, WaaF and Vpar 0760), GT28 (MurG), GT30 (WaaA and KdtA), GT35 (AtPHS2, MalP and GlgP), GT68 (POFUT2), GT80 (Pdst and Bst), GT4 (PimA, PimB, WsaF, GtfA, TarM, PglH, MshA, BshA and WaaG), GT_cp_ and its homologs (GT_la_, GT_tl_, GT_mn_, GT_dm_, GT_cb_, GT_kg_, GT_al_, GT_bb_, GT_dr_, GT_pa_, and Agt). GTpa is the GT4 glycosyltransferase of *Pseudomonas* sp. PA14 (Supplementary Figure S1). (B) Multiple sequence alignment for GT_cp_ and its homologs using Clustal W program with manual adjusting: GT_cp_ from *Candidatus* Pelagibacter sp. HTCC7211 (WP_008545403.1); GT_la_ from *Labrenzia aggregata*(WP_040439323.1); GT_tl_ from *Thalassospira lucentensis* (WP_062950653.1); GT_mn_ from *Methylophaga nitratireducenticrescens* (WP_014706011.1); GT_dm_ from *Desulfobulbus mediterraneus* (WP_028584068.1); GT_cb_ from *Citromicrobium bathyomarinum* JL354 (WP_010239457.1); GT_kg_ from *Kordiimonas gwangyangensis* (WP_051078133.1); the characterized Agt from *A. tumefaciens* (atu2297) and MshA from *C. glutamicum* (WP_143854623.1). Red bars represent α-helical regions and blue arrows represent β-sheet from MshA (Protein Data Bank [PDB] entry 3C4Q). Residues interacting with UDP-sugar are indicated by the closed black circles. Catalytic dyad composed of His104-Asp256 is shaded in grey. Black boxed regions and red boxed regions in the alignment indicated two conserved UDP-sugar binding motifs of GRVAXEKN and FPSXTDTFG and a conserved Gly-rich motif, respectively. (C) Homology modelling showing the predicted structure of GT_cp_ and the catalytic dyad composed of His104-Asp256. The signature glutamate residue in MshA (Glu316) is substituted to an aspartate residue (Asp256) in GT_cp_.

Due to the lack of a three-dimensional structure of DAG-dependent GT4 glycosyltransferases to date, a model of GT_cp_ was generated by homology modelling using the X-ray structure of the GT MshA co-crystalized with UDP (Protein Data Bank [PDB] entry 3C4Q; 17% sequence identity) from *Corynebacterium glutamicum* as the template (Fig. 1C) (22). MshA, also a member of the GT4 family, catalyses the first step of the biosynthesis of mycothiol in actinobacteria using UDP-*N*-acetylglucosamine (UDP-GlcNAc) as the sugar donor (22). Using the PDBeFold server, we compared the crystal structure of MshA with the modelled structure of GT_cp_. The predicted structure of GT_cp_ includes a GT-B fold consisting of two Rossmann-like *β-α-β* domains, with the N-terminal domain (residues 1–160 and residues 320–332) and C-terminal domain (residues 169–315), separated by a large cleft that includes the catalytic centre. The same GT-B fold is also found in several members of the GT4 family enzymes, including MshA and PimA (Fig. 1A) (22, 23). The substrate binding site of the sugar-donor is located mainly in the C-terminal domain, where the sugar-donor forms a number of hydrogen bonds with the protein. Given that the two enzymes (*i.e.* MshA and GT_cp_) share only 17% overall sequence identity, the alignment was manually corrected by incorporating information such as predicted secondary structures and conserved functional residues. Multiple sequence alignment of GT_cp_ and its homologs revealed that GT_cp_ contains a catalytic dyad composed of His104–Asp256, two conserved UDP-sugar binding motifs (GRVAXEKN and FPSXTDTFG), and a conserved Gly-rich motif, all of which are commonly found in the GT4 family (Fig. 1B) (24).

### Cloning of putative gene encoding GT_cp_ and functional verification

The recombinant plasmid pET22b–GT_cp_ was constructed to purify the enzyme for determination of its catalytic properties. Soluble expression of His-tagged GT_cp_ was achieved in *E. coli* BL21 (DE3) by adding 0.5 mM isopropyl-β-D-thiogalactopyranoside (IPTG). The recombinant GT_cp_ was purified to homogeneity by Ni–NTA affinity followed by gel filtration chromatography using the Superdex 200 column. In SDS-PAGE analyses, the purified recombinant His-tagged GT_cp_ was visible as a major band with a calculated mass of 38 kDa (Fig. 2A). The process used to purify GT_cp_ is summarized in Table 1. The enzyme was purified 3.5-fold with a yield of 17.1% and a specific activity of 47.1 U/mg. The predicted molecular mass of the monomeric form of GT_cp_ is 38 kDa, but GT_cp_ eluted as a peak corresponding to a molecular mass of approximately 70 kDa in the gel filtration chromatography experiments (Fig. 2B). These results suggested that GT_cp_ forms a dimer in solution, consistent with the proposed mechanism that oligomerization is a major factor contributing to the biochemical function and enzymatic activity of GTs (24).

**Figure 2.**
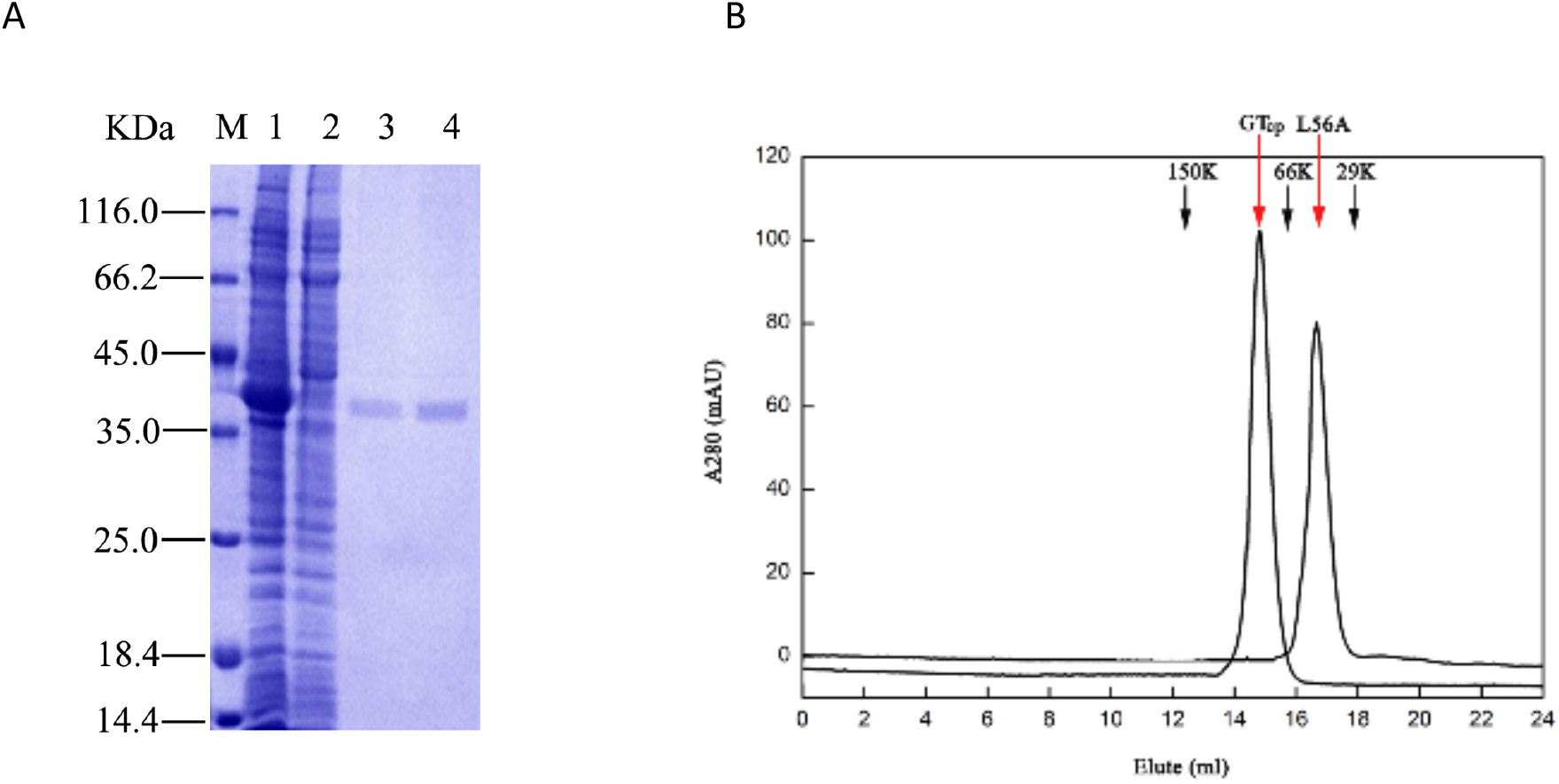
(A) Over-expression and purification of GT_cp_ proteins from *Candidatus* Pelagibacter sp. HTCC7211. M, protein molecular weight marker. Lane 1 cell-free supernatant induced with Isopropyl β-D-1-thiogalactopyranoside (IPTG), lane 2 cell-free supernatant without IPTG induction and lanes 3-4, purified GT_cp_ (molecular weight estimated to be ~ 38 kDa). (B) Gel filtration analysis of the wild-type GT_cp_ and the mutant L56A. The red arrows indicate the eluted position of GT_cp_ and the L56A mutant, the black arrows indicate protein markers (from left to right): alcohol dehydrogenase (150 kDa, 12.35 ml), albumin (66 kDa, 15.83 ml) and carbonic anhydrase (29 kDa, 17.95 ml).

**Table 1.** Purification of the recombinant GT_cp_ from *Candidatus* Pelagibacter sp. HTCC7211

| Step | Total protein<br>(mg) | Total<br>activity (U) | Specific<br>activity (U/mg) | Purification<br>fold | Yield (%) |
| --- | --- | --- | --- | --- | --- |
| Crude cell extract | 34.8 | 469 | 13.4 | 1 | 100 |
| Ni-NTA affinity | 1.9 | 85 | 44.7 | 3.3 | 18.1 |
| Superdex-200 gel filtration | 1.7 | 80 | 47.1 | 3.5 | 17.1 |

We used two methods to determine the activity of purified GT_cp_. Previous studies have shown that thin-layer chromatography (TLC) can resolve these selected glycoglycerolipid standards (15, 16). Indeed, as shown in Fig. 3A, the products of the enzymatic reaction with MGlc-DAG, MGal-DAG, and MGlcA-DAG were observed by staining with sulfuric acid/methanol/water, and corresponded to the standard markers. These results showed that GT_cp_ is able to transfer galactose, glucose, and hexuronic acid to the DAG acceptor using the respective UDP-sugars.

**Figure 3.**
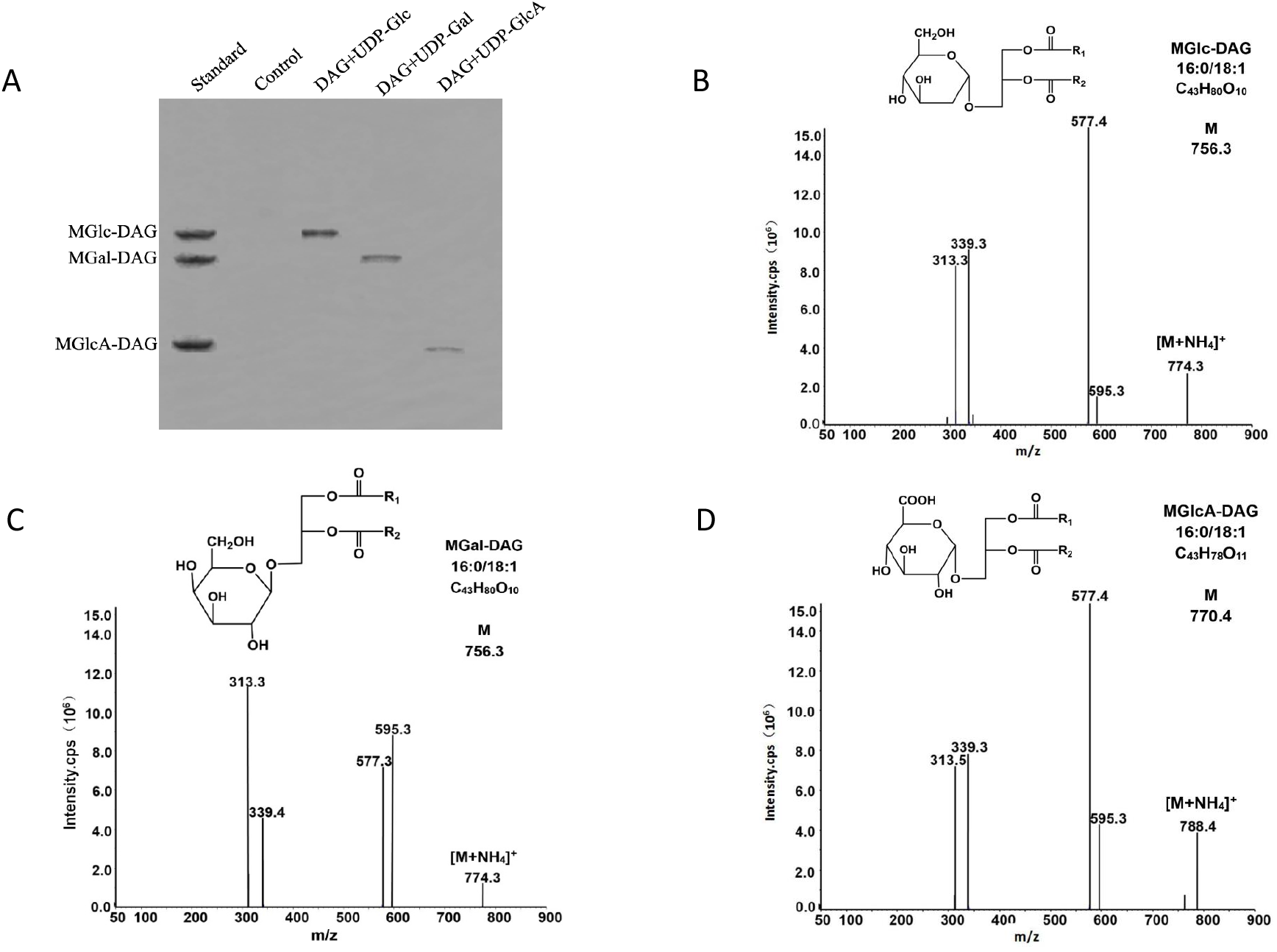
Functional characterization of recombinant GT_cp_. (A) TLC of the enzymatic reaction products with different UDP-sugar donors and diacylglycerol (DAG) as the acceptor by staining with sulfuric acid/methanol/water (45:45:10). (B-D) LC-MS of fragmentation spectra for monohexuronosyl DAGs (MGlc-DAG and MGal-DAG) and MGlcA-DAG were obtained from the products of the GT_cp_-catalysed reaction.

For structural identification, the glycoglycerolipids were analysed by liquid chromatography-mass spectrometry (LC-MS). LC-MS analyses (in the positive ion mode) detected an ammonium adduct (Fig. 3B-D) and fragmentation spectra for monohexuronosyl DAGs (MGlc-DAG and MGal-DAG) and MGlcA-DAG were obtained from the products of the GT_cp_-catalysed reactions. The calculated m/z of the parental ion of MGlc-DAG, MGal-DAG, and MGlcA-DAG was 756.3, 756.3, and 770.4, respectively. The two species differed in the neutral loss corresponding to the polar head group (179.0 and 193.1 m/z for the loss of the hexosyl group and the hexuronic acid group, respectively). In each case, this loss yielded a DAG-16:0/18:1 (m/z 577.3 or 577.4). A further two peaks corresponded to monoacylglycerol with glyceryl-16:0 (m/z 313.3 or 313.5) and −18:1 (m/z 339.3 or 339.4) fatty acids, respectively. These results demonstrated that GT_cp_ shows high enzymatic activity towards the synthesis of MGlc-DAG, MGal-DAG, and MGlcA-DAG from DAG and UDP-sugars. Several bacterial lipid GTs from *M. loti, A. tumefaciens, Mycoplasma pneumonia*, and *Mycoplasma genitalum* have been found to synthesize different glycoglycerolipids (DGal-DAG, GlcGal-DAG, and triglycosyl DAGs) using UDP-Glu and UDP-galactose (UDP-Gal) as sugar donors (15, 16, 25, 26). The *A. tumefaciens* GT Agt, which synthesizes neutral glycoglycerolipid (MGlc-DAG) and acidic glycoglycerolipid (MGlcA-DAG) with UDP-Glc or UDP-GlcA as the sugar donor, respectively, has been isolated and characterized (18). To the best of our knowledge, GT_cp_ is the first bacterial lipid GT acting as a multifunctional enzyme to synthesize MGlc-DAG, MGal-DAG, and MGlcA-DAG with different UDP-sugars as donors. It remains to be seen whether bi-functional/multifunctional GT4 enzymes involved in glycoglycerolipid synthesis are a common trait in this group. At least one member of this family, the GT4 homologue in *Pseudomonas* sp. appears to be specific for UDP-glucose and does not accept UDP-galactose nor UDP-glucuronic acid as the substrate (Supplementary Figure S1).

### Catalytic properties of GT_cp_

The catalytic activity of GT_cp_ was tested with UDP-Glc as the sugar donor substrate and DAG as the potential acceptor substrate. The effect of temperature on GT_cp_ activity was determined in the range of 10–50°C (Fig. 4A). GT_cp_ showed maximum activity at around 35°C and more than half of maximum activity at 20–45°C. Calculation of the activation energy *E_a._* using the Arrhenius plot in a semi-logarithmic form (ln *v→1/T*) was shown in Fig. 4B, and the slope of the plot was used to calculate the activation energy which was *E_a_* = 25.1 kJ mol^−1^. The thermostability of GT_cp_ was evaluated at three different temperatures (30°C, 40°C, and 50°C) with increasing incubation times up to 120 min. Most of the enzyme activity was maintained after incubation at 40°C for at least 120 min, whereas incubation at 50°C for 30 min reduced activity by approximately 50% (Fig. 4C). To investigate the effect of pH on the enzymatic activity of GT_cp_, the enzymatic reaction was evaluated in different buffers (pH 7.0–11.0). The maximum activity of GT_cp_ was at pH 8.5, and it retained more than 50% of maximum activity between pH 7.5 and 9.0 (Fig. 4D). Considering the significance of NaCl for marine enzymes, enzyme activity was determined in the presence of NaCl at different concentrations. The recombinant GT_cp_ maintained 54% of its maximum activity in the presence of 1.5 M NaCl and 30% of its maximum activity when the NaCl concentration was increased to 4 M (Fig. 4E).

**Figure 4.**
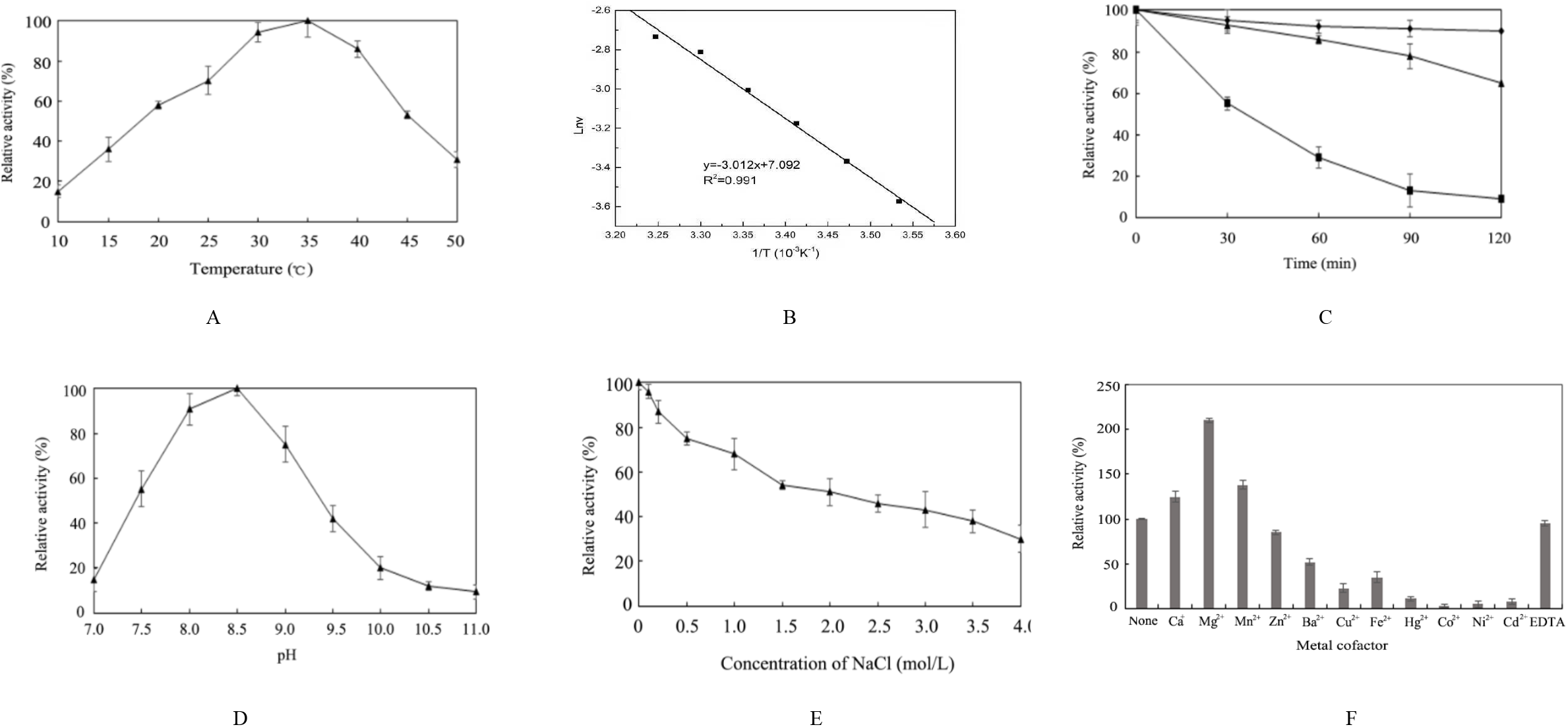
Biochemical characterization of GT_cp_. (A) Effect of temperature on the activities of GT_cp_. (B) Arrhenius plot of GT_cp_. The activation energy of the reaction *E_a_* = 25.1 kJ mol^−1^ could be determined from the slope of the regression curve. (C) Thermostability of GT_cp_. The residual enzyme activity was measured after incubation of the purified enzyme at 30°C (diamonds), 40°C (triangles), and 50°C (boxes), respectively. (D) Effect of pH on the activities of GT_cp_. (E) Effect of NaCl on the activities of GT_cp_. The enzyme was incubated in buffers containing different concentrations of NaCl (0 to 4 M) at 4°C for 1 h. Residual activity was measured under optimal conditions. F) Effect of metal ions (5 mM) on the activities of GT_cp_. The values are means of three independent experiments.

To investigate the substrate specificity of GT_cp_ for DAGs, different species of varying chain length of DAGs were tested (Table 2). The *k_m_* and *k_cat_* values of GT_cp_ were calculated from Hanes-Wolff plots and the Michaelis-Menten equation. For the DAGs with saturated fatty acid chains (di8:0, di10:0, di12:0, di14:0, di16:0, and di18:0), the *k_m_, k_cat_*, and *k_cat_/k_m_* values increased with increasing acyl chain length. GT_cp_ showed the higher activities for the unsaturated DAGs than for the saturated DAGs. The most preferred DAG substrate for GT_cp_ was C16:0/C18:1 DAG, consistent with the fact that C16:0 and C18:1 fatty acids are common in marine bacteria (17, 18).

**Table 2.** Kinetic parameters of GT_cp_ using UDP-Glc, UDP-Gal and UDP-GlcA as sugar donors and different DAG as acceptors ^a^.

| Substrate | $k_{cat}$ (min<sup>-1</sup>) | $K_m$ (μM) | $k_{cat}/K_m$ (min<sup>-1</sup> mM<sup>-1</sup>) |
| --- | --- | --- | --- |
| UDP-Glc/ DAG (C16:0 /C18:1) | 5.9±0.4 | 82.0±0.3 | 71.9±2.7 |
| UDP-Gal/ DAG (C16:0 /C18:1) | 1.0±0.3 | 32.0±0.4 | 31.3±2.9 |
| UDP-GlcA/ DAG (C16:0 /C18:1) | 4.1±0.1 | 68.0±0.5 | 60.3±3.6 |
| UDP-Glc/DAG (C18:0 /C20:4) | 4.8±0.2 | 77.1±0.4 | 62.2±0.9 |
| UDP-Glc/DAG (C18:0 /C18:2) | 6.2±1.5 | 102.5±0.8 | 60.5±1.8 |
| UDP-Glc/DAG (di18:1) | 5.0±0.8 | 90.5±1.6 | 55.2±2.2 |
| UDP-Glc/DAG (di18:0) | 3.6±0.7 | 75.3±1.2 | 47.8±1.3 |
| UDP-Glc/DAG (di16:0) | 1.7±1.4 | 55.2±0.9 | 30.8±2.5 |
| UDP-Glc/DAG (di14:0) | 1.0±2.2 | 45.9±1.1 | 21.7±2.8 |
| UDP-Glc/DAG (di12:0) | 0.5±0.4 | 21.6±0.8 | 13.9±1.7 |
| UDP-Glc/DAG (di10:0) | ND | ND | ND |
| UDP-Glc/DAG (di8:0) | ND | ND | ND |
| UDP-Glc/DAG (C16:0 /C18:1)<sup>b</sup> | 8.3±0.3 | 58.0±0.6 | 143.1±3.9 |

<sup>a</sup>The values are means of three independent experiments ± standard deviations. ND, not detectable.
<sup>b</sup>The kinetic parameters were determined using purified GT<sub>cp</sub> in the presence of Mg<sup>2+</sup> (5 mM).

The Michaelis-Menten kinetic parameters for GT_cp_ were determined using UDP-Glc, UDP-Gal, and UDP-GlcA as the sugar donors (Table 3). The *K*_m_ value for UDP-Glc (82 μM) was higher than those for UDP-Gal and UDP-GlcA, consistent with the fact that UDP-Glc is the preferred substrate at physiological condition (27). The *K*_cat_/*K*_m_ value for GT_cp_ toward different sugar donors followed the order UDP-Glc (71.4±2.7) >UDP-GlcA (59.6±3.6) >UDP-Gal (32.2±2.9). UDP-xylose, UDP-rhamnose, UDP-mannose and UDP-fructose were also tested with C16:0/C18:1 DAG, which showed no activity. Thus, GT_cp_ exhibited the highest enzymatic activity for UDP-Glc among the sugar donors tested. In contrast with GT_cp_, the Pgts from *M. loti* and *A. tumefaciens* favour uridine UDP-Gal over UDP-Glc (15, 16). A comparison of GT_cp_ kinetics with other biochemically characterized GT4 glycosyltransferases is listed in Table 4 (22, 28–33).

**Table 3.** Kinetic parameters of GT_cp_ using UDP-Glc, UDP-Gal and UDP-GlcA as sugar donor and C16:0/C18:1 DAG as acceptor ^a^.

| Substrate | $k_{cat}$ (min<sup>-1</sup>) | $K_m$ (μM) | $k_{cat}/K_m$ (min<sup>-1</sup> mM<sup>-1</sup>) |
| --- | --- | --- | --- |
| UDP-Glc | 5.85±0.4 | 82±0.3 | 71.4±2.7 |
| UDP-Gal | 1.03±0.3 | 32±0.4 | 32.2±2.9 |
| UDP-GlcA | 4.05±0.1 | 68±0.5 | 59.6±3.6 |
| UDP-Glc (with 5 mM Mg<sup>2+</sup>) | 8.25±0.3 | 58±0.6 | 142.2±3.9 |

<sup>a</sup>The values are means of three independent experiments.

**Table 4.** Kinetic parameters of GT_cp_ in comparison with GT4 glycosyltransferases using different sugar donor ^a^.

| Parameter | GT<sub>cp</sub> | PimA | PimB | MshA | TarM | BshA | GtfA | PglH |
| --- | --- | --- | --- | --- | --- | --- | --- | --- |
| Donor substrate | UDP-Glc | GDP-Man |  | UDP-GlcNAC |  |  |  |  |
| $K_m$ ($\mu M$) | 82 ± 0.3 | 18 ± 2 | 19.0 ± 4.6 | 0.208 ± 0.017 | 65 ± 10 | 180 ± 50 | 11.8 ± 1.5 | 2.6 ± 0.3 |
| $k_{cat}$ ($min^{-1}$) | 5.8 ± 0.4 | | | | 126 ± 10 | 78.6 | 7.35 ± 0.42 | 4.4 ± 0.2 |
| $k_{cat}/K_m$ ($min^{-1}mM^{-1}$) | 71.4 ± 2.7 | | | | | 6.8 | 0.63 | 1.7 ± 0.2 |

<sup>a</sup> The values are means of three independent experiments. UDP-GlcNAC: UDP-N-acetylglucosamine; GDP-Man: GDP-mannose

### Metal ions improve enzyme activity of GT_cp_

The effect of various metal ions on the enzyme activity of GT_cp_ is shown in Fig. 4F. Among the tested metal ions (5 mM), Mg^2+^, Ca^2+^, and Mn^2+^ significantly stimulated GT_cp_ activity by up to 223%, 125%, and 138%, respectively, although the as-isolated enzyme is already active Furthermore, EDTA did not significantly affect the enzymatic activity of GT_cp_ after incubation for 60 min at room temperature. The activity of GT_cp_ was decreased by Ba^2+^, Cu^2+^, and Fe^2+^ to 52%, 23%, and 35%, respectively. Moreover, Hg^2+^, Cd^2+^, Ni^2+^, and Co^2+^ completely abolished enzyme activity. This may have resulted from the binding of these metal ions to the -SH, -CO, and -NH moieties of the amino acids of GT_cp_, leading to structural changes and inactivation (34).

Given that Mg^2+^ markedly improved the activity of GT_cp_, we determined metal content of the purified enzyme by inductively coupled plasma-mass spectrometry (ICP-MS) and found that Mg^2+^: protein molar ratio was < 0.03 (supplementary table 1). Similarly, no substation amount of Ca, Mn or Zn was found in GT_cp_, suggesting that this enzyme is unlikely a metalloprotein. The kinetic parameters were subsequently determined using purified GT_cp_ in the presence and absence of Mg^2+^ (Table 3). In the presence of excess DAG, the *K*_m_ value of GT_cp_ decreased from 82 μM to 58 μM with the addition of Mg^2+^, indicating that GT_cp_ has a higher affinity for UDP-Glu in the presence of Mg^2+^. The catalytic efficiency was nearly 2.0-fold higher than that in the absence of Mg^2+^. In the presence of excess UDP-Glc, the *K*_m_ was almost unchanged with/without Mg^2+^. Metal ions are important regulators of physiological functions and contribute to the preservation of the structural integrity of some proteins (35). In contrast to GT-A fold GTs, GT-B fold GTs, including GT_cp_, are metal ion-independent (24). However, some studies have found that metal ions also change GT-B fold activity, such as GGT58A1 from *Absidia coerulea* (36), UGT59A1 from *Rhizopus japonicas*, Bs-PUGT from *Bacillus subtilis* P118 (37), and human POFUT2 (38). In some cases, metal ion simultaneously interacts with both the enzyme and the sugar donor in the active site, which causes the glucosyl donor to realign in the active site, and thus affects the activity of the enzyme (39). Interestingly, although human POFUT2 is not a metalloprotein, Mg^2+^, Mn^2+^ and Ca^2+^ are also known to enhance the enzyme activity by facilitating the release of product from the enzyme (38). Therefore, it is tempting to speculate that the addition of Mg^2+^ may also help enhance GT_cp_ catalysis in a similar manner.

### Insights into structure and glycosylation mechanism of GT_cp_

GT_cp_, along with the previously characterized Agt, form a group in the GT4 family and are GT-B fold GTs (Fig. 1, Fig. 5A). The GT-B fold GTs are thought to employ the so-called retaining glycosylation for adding the sugar unit to the substrate although some researchers have proposed an alternative internal return (SN*i*-like) mechanism (24, 40). In the latter model, the nucleophilic attack and the departure of the leaving group occur on the same face of the sugar, and involve the formation of a short-lived oxocarbenium-like transition state with asynchronous acceptor glycoside bond formation and phosphate bond breakdown (41–43). According to this latter model, two conserved amino acid residues are important for catalysis (*e.g.* Glu316 and His109 in MshA). In MshA, Glu316 and His109 act as catalytic nucleophiles in the SN*i*-like mechanism, and form hydrogen bonds with the OH-3 and OH-6 of the glucosyl moiety, respectively (22). Indeed, we also found the corresponding residues (Asp256, His104) in GT_cp_ (Fig. 1B). Asp256 and His104 fulfil a critical role in GT_cp_ catalysis, because their substitution to Ala completely abolished catalytic activity (Fig. 5B). Interestingly, mutation of the conserved Asp residue at position 256 to Glu (D256E) reduced activity by more than 90% even though both Asp and Glu residues have a carboxylic acid moiety. Asp256 is essential for the enzymatic activity of GT_cp_ and cannot be replaced by Glu, indicating that the length of the side chain of this residue is important for the activity of GT_cp_. These results suggested that Asp256 and His104 play important functions in glycosyl transfer while maintaining the α-configuration of the anomeric carbon. To confirm the glycosylated position and the anomeric stereoselectivity, MGlc-DAG was purified and analysed by ^1^H-nuclear magnetic resonance (NMR) (Fig. 5C). The glucosyl moiety of the glycolipid was suggested by the characteristic signals for Glc-H1 (δH 5.17, 1H) and Glc-H2–6 (δH 3.20 to 3.70, 6H) by ^1^H NMR. The location of the glucosyl moiety was indicated by the correlation between Glc-H1 and C-3 (δH 136.0) in the heteronuclear multiple bond correlation (HMBC) spectrum. The large coupling constant (J=3.7Hz) of Glc-H1 revealed the α-D-configuration of the glucosidic linkage.

**Figure 5.**
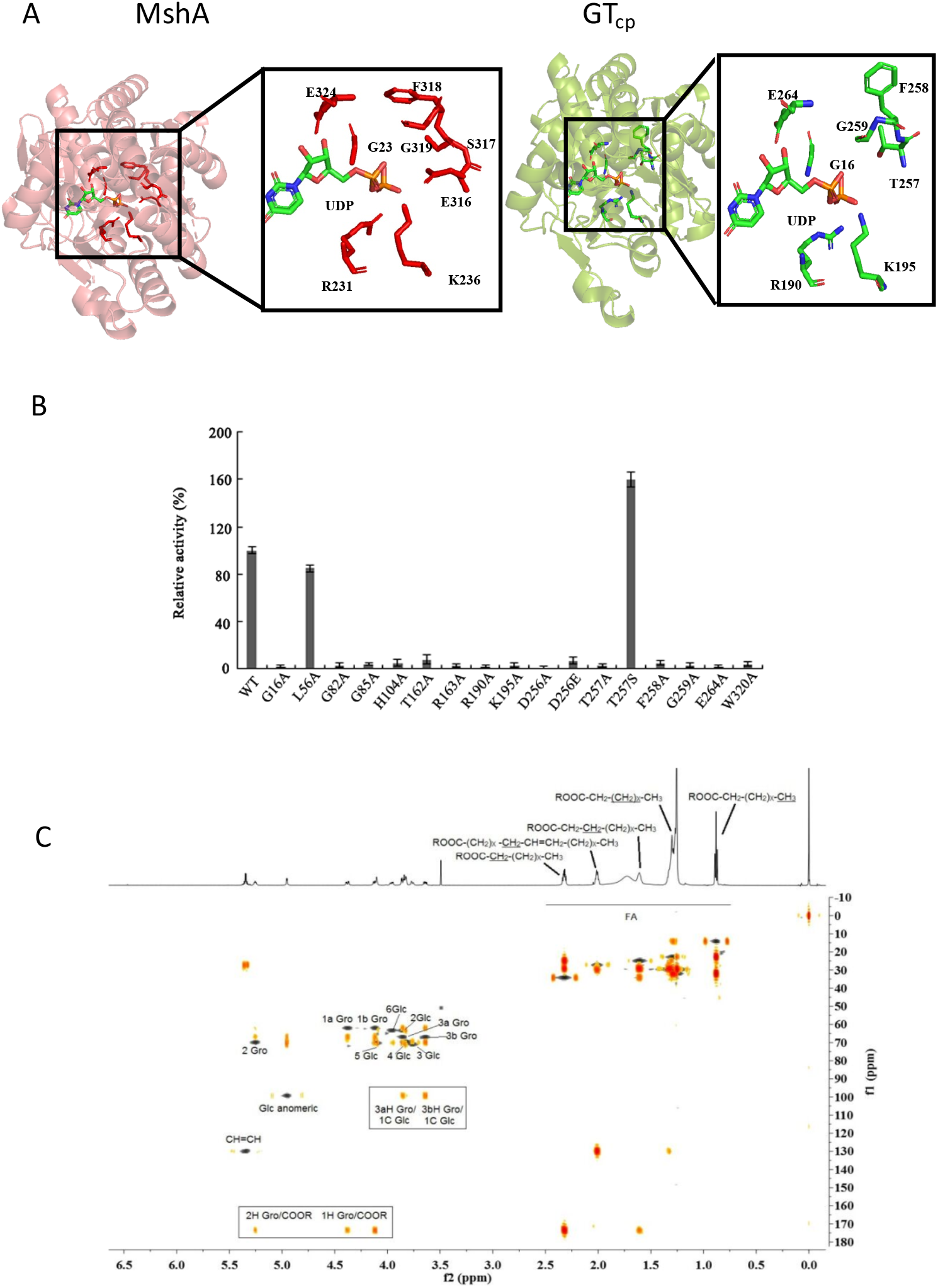
Homology modelling prediction of the UDP-sugar donor binding pocket in the GT_cp_. (A) UDP-sugar binding site of MshA (22) and predicted UDP-sugar binding site of GT_cp_. (B) Mutational analysis of the key amino acids involved in the catalytic glycosylation reactions of GT_cp_. C) Overlay of ^1^H, heteronuclear single-quantum correlation (HSQC) and heteronuclear multiple bond correlation (HMBC) spectra of MGIc-DAG.

In terms of the donor binding pocket, GT4 family GTs have two highly conserved sequence motifs in their C-terminal domain that are involved in the binding of the sugar nucleotide with the UDP moiety (23, 29, 44). The sequence alignment analyses indicated that GT_cp_ and its homologs have these two conserved motifs in the donor binding pocket (Fig. 1B), consistent with other GT4 proteins. The highly conserved sites were further investigated to illuminate the mechanism of sugar donor binding (Fig. 5A). Each of the potential active sites in the donor binding pocket (Gly16, Gly82, Gly85, Arg190, Lys195, Thr257, Phe258, Gly259, and Glu264) was mutated to Ala. Enzymatic activity of GT_cp_ in all the mutants was almost completely abolished, supporting our assumption from the homology model that these residues are integral to the sugar donor binding site (Fig. 5B). Thr257 in the structure of GT_cp_ corresponds to MshA residue Ser317, which interacts with the 4-OH of the sugar moiety and forms hydrogen bonds (22). When the residue at position 257 (Thr) in GT_cp_ was substituted with Ser (T257S), the T257S mutant exhibited approximately 1.6-times higher activity than that of wild-type GT_cp_. Compared with the parental residue Thr, Ser has one fewer methyl group; therefore, there is more free space between the Ser residue and the substrate (45). The generated space might allow more room for the appropriate interaction between the Ser residue and the sugar donor. These findings indicate that the mechanism of the sugar donor recognition of GT_cp_ is similar to that of known GT-B fold GTs, such as MshA. That is, the conserved key residues would form hydrogen bonds with the UDP part and interact with the sugar moiety of the sugar donor.

In order to identify potential residues involved in binding of DAG, we docked C16:1/C18:0 DAG into the homology model of GT_cp_. DAG is predicted to lay in the groove of the open structure in the model (Fig. 6A, B). Docked DAG interacts with several key resides primarily through hydrophobic interactions but also some polar interactions *e.g.* His104, Thr162 and Arg163 and Trp320 (Fig. 6C), of which His104, Arg163 and Trp320 are strictly conserved in GT_cp_ in a range of bacteria (Fig. 1B). Indeed, the His104, Arg163 and Trp320 mutants are inactive, supporting a key role in GT_cp_ catalysis (Fig. 5B). Subsequent independent docking of the three UDP-sugars predicted the identical binding pocket for all three (Fig.6D) in the presence of already docked DAG and with UDP-galactose as an example we observe the DAG and UDP-sugars positioned parallel to one another in the ‘open state’ (Fig. 6 E). To the best of our knowledge, our study provides the first insight of the binding pocket of DAG in a GT4 glycosyltransferase involved in glycoglycerolipid biosynthesis.

**Figure 6.**
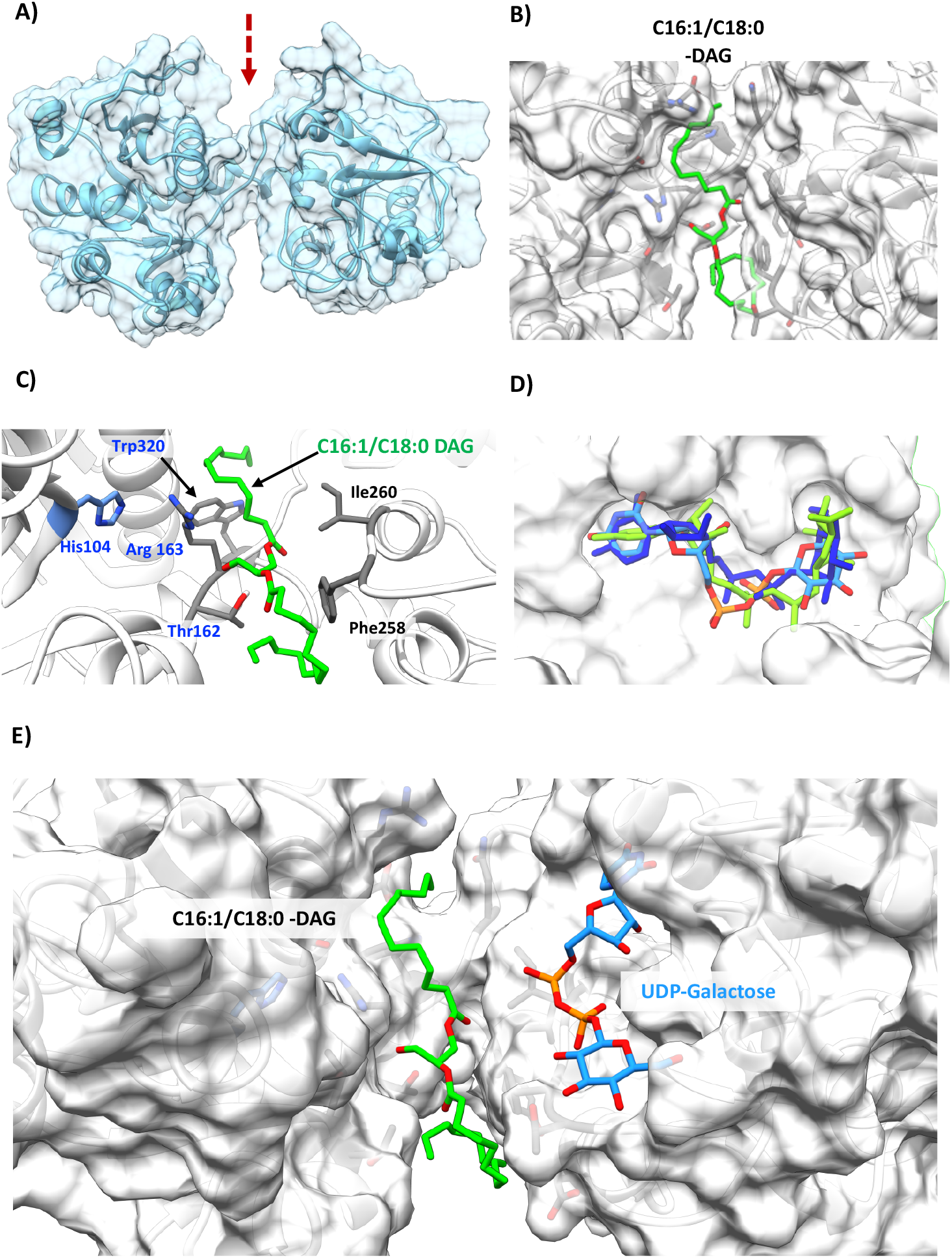
Identification of key residues for diacylglycerol (DAG) binding in the GT_cp_. **A)** Homology modelling prediction of the open model based on template 2R60. The arrow indicates the wide cleft. **B)** C16:1/C18:0-DAG (green) docked in GT_cp_ in the groove (shown as a transparent surface). **C)** A detailed depiction of key coordinating hydrophobic (black) and polar (light blue) residues for DAG. His104, Thr162, Arg163, Trp320 are crucial for enzyme activity (Fig. 5B). **D)** An overlay of UDP-glucose (Blue), UDP-galactose (Cyan) and UDP-glucuronic acid (Green) showing all three ligands can occupy the same binding site in similar poses in the DAG-docked GT_cp_. **E)** A top-down view of DAG and UDP-galactose shown docked parallel to each other in their respective binding pockets/grooves.

In MshA, the hydrophobic residue Leu76 helps to stabilize the dimer (22). This residue corresponds to Leu56 in GT_cp_ (Fig 1B). To investigate the role of Leu56 in GT_cp_, a site-directed mutant to Ala was made. The mutant L56A protein was loaded onto a Superdex 200 column to analyse its oligomeric state. The eluted peaks of L56A mutant corresponded to the monomer of GT_cp_ (38 kDa) according to their elution volumes (Fig. 2B). No difference was observed between the wild-type profile and the other mutants during protein purification (data not shown). The mutant L56A showed approximately 85% of wild-type GT_cp_ activity. Structural studies revealed that most of the residues involved in oligomerization are conserved with related GTs, and they appear to be primarily hydrophobic and aromatic residues that form an extensive hydrophobic interface between the monomers (46). These results demonstrated that residue Leu56 is essential for the stable dimerization of the protein, but does not play a direct role in the catalytic reaction of GT_cp_.

To conclude, our data show that the activity of purified GT_cp_ from the marine bacterium *Candidatus* Pelagibacter sp. HTCC7211 is sufficient for the synthesis of several glycoglycerolipids, including MGlc-DAG, MGal-DAG and MGlcA-DAG. The ability to synthesize MGlc-DAG and MGlcA-DAG suggest that GT_cp_ may play an important role in lipid remodelling in natural marine systems. GT_cp_ and PlcP, a manganese-dependent metallophosphoesterase, are organized in an operon-like structure in numerous marine heterotrophic bacteria (17, 18). Upon phosphorus (P) deficiency, PlcP selectively degrades phospholipids such as phosphatidylglycerol (PG) and phosphatidylethanolamine (PE) to DAG, which then serves as the substrate for the biosynthesis of these glycolipids by GT_cp_ using UDP-Glc, UDP-Gal, or UDP-GlcA as sugar donors (Fig. 7). Both the phospholipid PG and the glycoglycerolipid MGlc-DAG are anionic under physiological conditions and it is likely that they could be interchangeable while maintaining the desirable biophysical properties of the membrane. Indeed, this has also been documented in the SAR11 strain HTCC7211 (17, 18). Similarly, substitution of PG by the anionic sulfur-containing glycolipid SQDG has also been shown for marine cyanobacteria and phytoplankton (47). Together, our work thus points to the important role of glycosyltransferases as key enzymes in the synthesis of glycoglycerolipid in marine bacteria.

**Figure 7.**
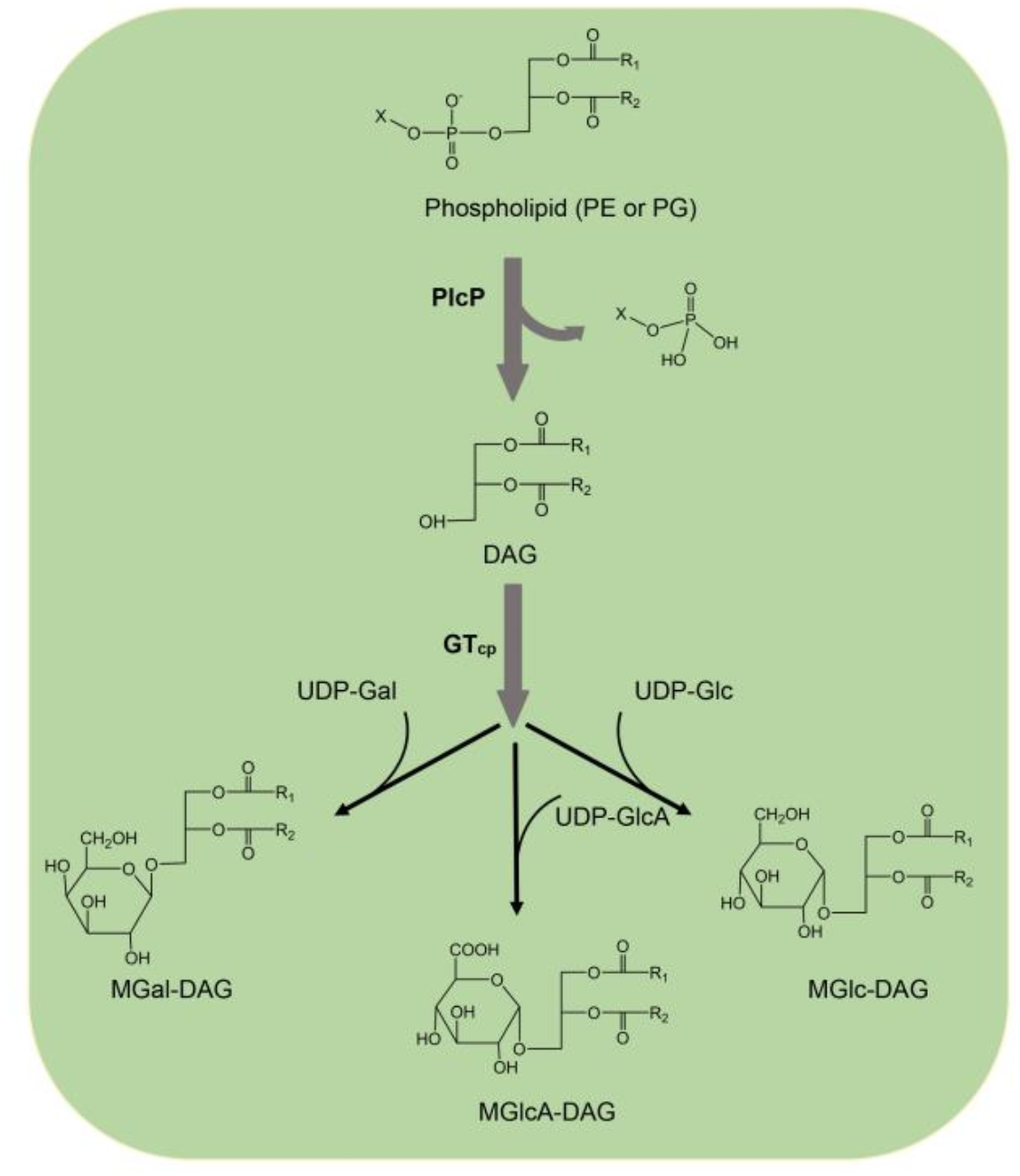
Proposed pathway of the synthesis of non-phosphorus glycoglycerolipids through PlcP and GT_cp_ in *Candidatus* Pelagibacter sp. HTCC7211. PlcP converts phosphatidylglycerol (PG) or phosphatidylethanolamine (PE) to generate diacylglycerol (DAG), and GT_cp_ can synthesize different glycoglycerolipids MGlc-DAG, MGal-DAG and MGlcA-DAG with UDP-Glc, UDP-Gal, or UDP-GlcA as sugar donors and DAG as the acceptor.

## Materials and Methods

### General materials and microorganisms

We purchased UDP-glucose (UDP-Glc), UDP-galactose (UDP-Gal), UDP-glucuronic acid (UDP-GlcA), UDP-xylose, UDP-rhamnose, UDP-mannose, UDP-fructose and diacylglycerols (DAGs) from Sigma-Aldrich Co. (St. Louis, MO, USA). Nickel column and Superdex 200 gel filtration columns were from GE Healthcare (Buckinghamshire, UK). All other chemicals were of the highest reagent grade and were obtained from Sangon (Shanghai, China). The *E. coli* strains JM109 for DNA manipulation and BL21-CodonPlus (DE3)-RIL for protein expression were obtained, respectively, from TaKaRa Bio, Inc. (Dalian, China) and Stratagene (La Jolla, CA, USA).

### Cloning, expression, and purification of GT_cp_

The gene *GT_cp_* from *Candidatus* Pelagibacter sp. HTCC7211 was codon-optimized and chemically synthesized by Sangon (Shanghai, China). Several site-directed *GT_cp_* mutants (encoding mutations G16A, L56A, G82A, G85A, H104A, T162A, R163A, R190A, K195A, D256A, D256E, T257A, T257S, F258A, G259A, E264A and W320A) were constructed using the overlapping PCR method with the common primers as for the wild-type GT_cp_ and two site-specific primers for each mutant (Table 5). The genes were then cloned into the pET22b expression vector using *Nco*I and *Sal*I restriction sites, and transformed into the host *E. coli* BL21-CodonPlus (DE3)-RIL for gene expression. The transformed cells were grown in LB medium containing 100 mg/l ampicillin and 34 mg/l chloramphenicol at 37°C with shaking at 180 rpm. When the cultures reached an OD 600 of 0.6, IPTG was added to a final concentration of 0.5 mM. After a further 4 h of growth at 37°C, the cells were harvested by centrifugation and lyophilized by vacuum-freezing. The harvested cells were re-suspended in buffer A (50 mM Tris–HCl, pH 7.9, 50 mM NaCl), with 1% (w/v) of triton X-100, and then disrupted by sonication. The cell mixture was then centrifuged at 12,000×g for 30 min, and the soluble fraction was loaded onto a nickel column (GE Healthcare) pre-equilibrated with buffer A. The recombinant enzymes were eluted with elution buffer (20 mM Tris–HCl, pH 7.9, 500 mM NaCl, 300 mM imidazole) and dialyzed overnight in buffer A to remove imidazole. For further purification, the enzymes were loaded on a Superdex 200 (16/60) gel filtration column (GE Healthcare), which was pre-equilibrated with buffer B (50 mM Tris–HCl, pH 7.9, 200 mM NaCl). The fraction size was 0.5 ml and the flow rate was 0.5 ml/min. The peak fractions were collected, concentrated, and analysed by SDS-PAGE (12% polyacrylamide). The protein concentration was determined using the Bradford method. The purified protein was stored in buffer A containing 25% glycerol at −80°C. SDS-PAGE gels and circular dichroism (CD) spectroscopy analyses of the purified protein and the mutants are shown in supplementary Figure S2.

**Table 5.**
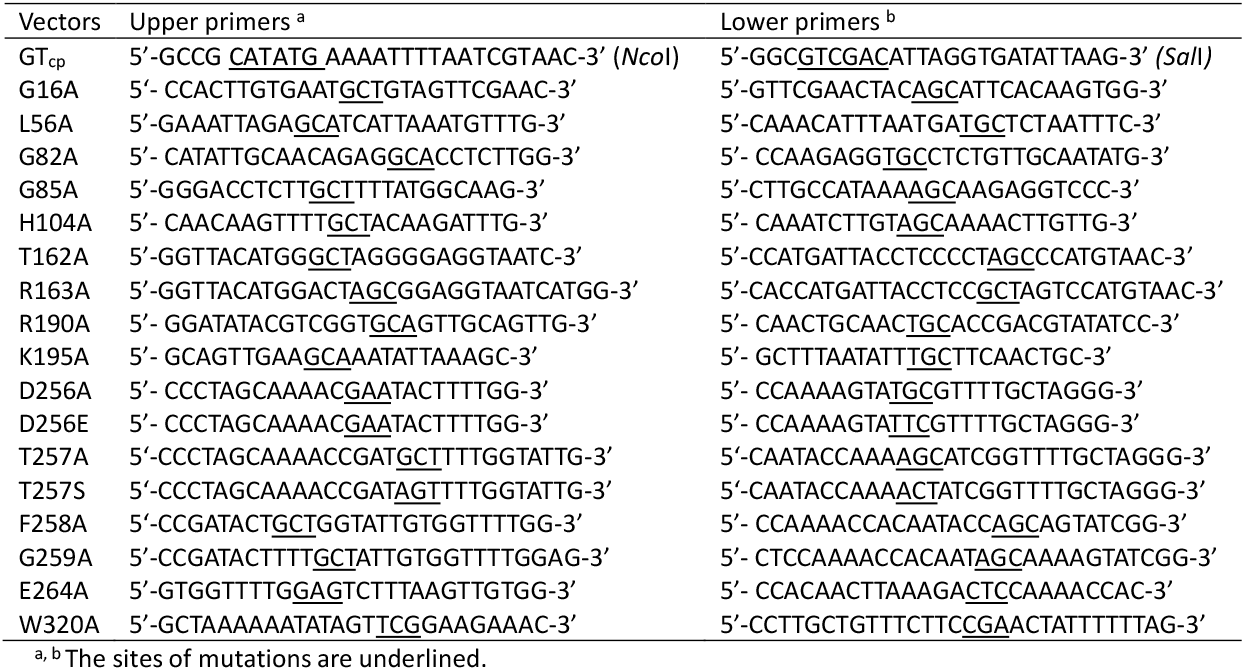
Primers used for PCR amplification in this study

### Bioinformatics and homology modelling

A putative GT gene encoding GT_cp_ was identified in the genome of *Candidatus* Pelagibacter sp. HTCC7211 (GenBank accession no. WP_008545403.1). ClustalW2 software (http://www.ebi.ac.uk/Tools/clustalw2/index.html) was used for multiple sequence alignment analysis of GT_cp_ (48). For the phylogenetic analysis, we used the neighbour-joining method and Molecular Evolutionary Genetic Analysis 7.1 software (MEGA, version 7.1) (49). The three-dimensional model structure of GT_cp_ was generated using tools at the Phyre 2 protein modelling server (50) and the crystal structure of MshA (Protein Data Bank [PDB] entry 3C4Q) from *C. glutamicum* as the template. Docking of UDP-Gla/UDP-Glc/UDP-GlcA with GT_cp_ was predicted using the Flexible Docking module in Accelrys Discovery studio (51, 52), and the protein model was imported into Flare (v3.0, Cresset) for docking the DAG substrate and firstly energy minimized with 2000 iterations with a cut off of 0.200 kcal/mol/A. The DAG lipid was imported as a ligand and energy minimized in Flare before being docked into the active site and the best scoring pose selected. Thereafter we utilised the DAG docked ligand with the model as the basis for the follow up *in silico* docking of UDP-sugars in the presence of the DAG.

### Enzyme activity assay

The enzymatic activity of GT_cp_ was measured using 0.1 mM UDP-Glc (or UDP-Gal, or UDP-GlcA) and 0.1 mM DAG as the substrate and 2.0 μM purified enzyme in 10 mM Tricine/KOH, pH 8.5, 2 mM DTT. The resulting mixture (500 μl) was incubated at 35°C for 60 min with constant shaking at 200 rpm. The products (glycoglycerolipids and DAGs) were extracted using the Floch method with methanol-chloroform-water at a ratio of 1:2:0.6 (vol/vol/vol). The lipid extract was dried under nitrogen as at room temperature. The dried lipids were resuspended in acetonitrile and ammonium acetate (10 mM, pH 9.2) at a ratio of 95:5 (vol/vol) and analysed by LC-MS. One unit of enzymatic activity was defined as the amount of enzyme required to catalyse the conversion of 1 μmol DAG per min under the standard conditions. The measurements were corrected for background hydrolysis in the absence of the enzyme. The *K*_m_ and *v*_max_ values were calculated using Hanes-Wolff plots with various concentrations of substrate (0.02 to 1.0 mM) and three replicates.

### Lipid analysis by TLC and LC-MS

The GT_cp_-synthesized glycoglycerolipids were analysed by TLC using a Camag Automatic TLC Sampler III (Camag, Muttenz, Switzerland) for spotting. Glycoglycerolipids were separated on silica gel 60 (Merck, Darmstadt, Germany) with chloroform/methanol/water (65:35:4 vol/vol), and stained with sulfuric acid/methanol/water (45:45:10 vol/vol) for visualization. The resulting solutions were further analysed by high-performance liquid chromatography using a 1290 Infinity II UPLC instrument (Agilent Corp., Santa Clara, CA, USA) coupled with an AB SCIEX Triple Quad™5500 (AB SCIEX, Framingham, MA, USA) equipped with an electrospray-ion (ESI) detector. A BEH Amide XP column (2.5-μm inner diameter, 3 mm by 150 mm, Waters, Milford, MA, USA) was used for chromatographic separation. The mobile phase consisted of acetonitrile (solvent A) and 10 mM ammonium acetate, pH 9.2 (solvent B). The column was equilibrated for 10 min with 95% A: 5% B prior to sample injection. The separation was conducted using a stepwise gradient starting from 95% A: 5% B to 70% A: 30% B after 15 min with a constant flow rate of 150 μl min^−1^. Mass spectrometric analysis was performed in the ESI positive ion mode with the ion spray voltage at 3500 V and temperature at 350°C. The nebulizer gas and heater gas were set at 40 psi. The analytical data were processed by Analyst software (version 1.6.3).

### Inductively coupled plasma-mass spectrometry (ICP-MS)

The metal content of GT_cp_ was measured by using an ICP-MS (Agilent Technologies 7900 ICP-MS). The standards for calibration were freshly prepared by diluting Ca, Mg, Mn, Zn and S stock solution (at 1000 mg·L^−1^ · Sigma-Aldrich, Saint Louis, MO, USA) with 1% (v/v) nitric acid with concentrations from 0.1 to 2.0 mg·L^−1^ for Ca, Mg, Mn, Zn, and from 1 to 25 mg·L^−1^ for S. About 3.0 mg protein was digested in 1% (v/v) nitric acid matrix for metal analyses. The content of S was quantified in order to determine the protein concentration. The contents of Ca, Mg, Mn, Zn and S were measured using the emission lines of 396.847 nm (Ca), 280.270 nm (Mg), 259.373 nm (Mn), 213.856 nm (Zn) and 180.669 nm (S) respectively.

### Nuclear magnetic resonance spectroscopy (NMR) spectroscopy

NMR spectroscopy experiments were carried out in CDCl3 with tetramethylsilane as an internal standard. ^1^H, ^13^C, DEPT, COSY, HSQC, HMBC and ROESY experiments were recorded at 298 K with 600 MHz spectrometer (Bruker Avance 600, Bruker). Bruker standard software Topspin 3.2 was applied to acquire and process all the spectra data. COSY and ROESY experiments were recorded using data sets (t1 by t2) of 2048 by 256 points, COSY with 4 and ROESY with 16 scans.

### Characterization of recombinant GT_cp_

The optimum temperature of GT_cp_ was measured in Tricine/KOH buffer (pH 8.5) in the range of 10–50 °C. The buffer was adjusted to pH 8.5 for each of the assayed temperatures. The activation energy of the cleavage reaction was calculated using the logarithmic form of the Arrhenius equation: ln*K_cat_*= ln*K_0_* -*E_a_*/R·T. The effect of pH on enzymatic activity was tested at 35°C for pH values in the range of 7.0–11.0. The following buffers were used: sodium phosphate (pH 6.0–7.5), Tricine/KOH (pH 7.5–9.5), and N-cyclohexyl-3-aminopropanesulfonic acid (CAPS, pH 9.5–11.0). The thermostability of the purified GT_cp_ was examined by incubating the enzyme in 50 mM Tricine/KOH buffer (pH 8.5) at three different temperatures (30, 40, and 50 °C). Samples (80 μL) of the enzyme were collected after incubation periods of 30, 60, 90, and 120 min at each temperature and the residual activity of each sample was assayed under standard conditions. The enzymatic activity of GT_cp_ was also measured in the presence of various metal salts (MnCl_2_, ZnCl_2_, MgCl_2_, CaCl_2_, BaCl_2_, CdCl_2_, HgCl_2_, CuCl_2_, FeSO_4_, NiSO_4_, and CoCl_2_) at 5 mM or 5 mM EDTA. To determine the salt stability of the enzyme, 0–4 M NaCl (final concentration) was added to the reaction mixture and enzyme activity was determined using optimum conditions.

## Data availability

The authors confirm that the data supporting the findings of this study are available within the article and its supplementary materials.

## Acknowledgements

This project has received funding from the European Research Council (ERC) under the European Union’s Horizon 2020 research and innovation programme (grant agreement no. 726116), Royal Society International Exchanges 2017 Cost Share (China) (lEC\NSFC\170213; grant agreement no. 170213), Program for Science &Technology Innovation Talents in the Universities of Henan Province (18HASTIT040) and Projects of science and technology activities for overseas students from department of human resources and social security of Henan Province (2019-3)

